# Phylogeography and population genetic structure of red muntjacs: Evidence of enigmatic Himalayan red muntjac from India

**DOI:** 10.1101/2020.08.11.247403

**Authors:** Bhim Singh, Ajit Kumar, Virendra Prasad Uniyal, Sandeep Kumar Gupta

## Abstract

We investigated the phylogeographic pattern of red muntjac across its distribution range, intending to address the presence of distinct lineages from Northwest India. The Complete mitogenome analysis revealed that India holds three mitochondrial lineages of red muntjac, whereas four were identified from its entire distribution range: Himalayan red muntjac (*M. (m.) aureus*), Northern red muntjac (*M. vaginalis*), Srilankan and Western Ghat India (*M. malabaricus*) and Southern red muntjac (*M. muntjak*) from Sundaland. The newly identified Himalayan red muntjac found in the Northwestern part of India, which was previously described based on their morphological differences. Estimates of the divergence dating indicate that the Northwest and Northern lineage split during the late Pleistocene approximately 0.83 Myr (CI_95%_:0.53 to 1.26), which is the younger lineages, whereas *M. malabaricus* is the most primitive lineage among all the red muntjac. Microsatellite results also supported the mitochondrial data and evident the presence of three distinct genetic clusters within India. The pronounced climate fluctuation during the Quaternary period was considered as a critical factor influencing the current spatial distribution of forest-dwelling species that restricted themselves in northwest areas. Based on molecular data, this study provided evidence of a new lineage within the red muntjac group from India that required to be managed as an evolutionary significant unit (ESUs). It highlighted a need for the taxonomic revision of Himalayan red muntjac (*M. (m.) aureus*) and also suggested its conservation status under IUCN Red List.

## INTRODUCTION

The genus *Muntiacus* belongs to the tribe Muntaicini and family Cervidae. It is widely distributed throughout South and Southeast Asia (Nagarkoti and Thapa, 2007) and exhibited dramatic variations in chromosome numbers (Yang et al., 1997; Wang and Lan, 2000). The taxonomic classification and validation of species and subspecies are still controversial. Mattioli, (2011) classified red muntjacs as single species *Muntiacus muntjak* that includes ten subspecies. Grubb and Groves (2011) recognized six species, and two subspecies of *Muntiacus* using geographical distributions and detail morphological characters. Recently, IUCN provisionally adopted two species of *Muntiacus: M. vaginalis* (Northern or Indian Red Muntjac) widely distributed from northern Pakistan to most of India, Nepal, Bhutan, Bangladesh to southern China including Hainan and south Tibet, and into Myanmar, Thailand, Lao, PDR, Viet Nam, Cambodia, whereas *M. muntjak* (Southern Red Muntjac) limited to Thai–Malay peninsula, Java, Bali, Lombok, Borneo, Bangka, Lampung and Sumatra and (Timmins et al., 2016). However, the exact southern range limit in the Thai–Malay peninsula remains unclear. For highly adapted mammals such as red muntjac whose extensive distribution ranges are linked to different political boundaries, contemporary genetic variation and population structure may be shaped by both natural and anthropogenic factors (Hewitt, 2000). As the recent studies on muntjac are increasing, the numbers of new species of *Muntiacus* are also increased, for examples based on the difference in skin color, skull and antler of museum specimens, a new endemic species from Borneo was described, the Bornean yellow muntjac (*M. atherod*) (Groves and Grubb, 1982) after that the Gongshan muntjac (*M. gongshanensis*) was described from Yunnan, China (Ma and Wang, 1990). Based on molecular studies, the Putao muntjac (*M. putaoensis*) from Myanmar (Amato et al., 1999), small blackish muntjac (*M. truongsonensis*) from central Vietnam (Giao et al., 1998), Roosevelt’s barking deer (*M. rooseveltorum*) from Vietnam (Minh et al., 2014) has already been discovered. Recently, complete mitogenome sequences reveal the presence of three distinct maternal lineal groups across the distribution ranges of *Muntiacus*: Srilankan red muntjacs that is confined to the Western Ghats, India and Sri Lanka; northern red muntjacs distributed in north India and Indochina; and southern red muntjacs found in Sundaland (Martin et al., 2017). Moreover, distinct morphological and genetic characters are drawing a center of attention for the researcher to identify more lineages of mystifying red muntjacs. Especially when human-mediated effects and several anthropogenic activities, such as habitat fragmentations/destruction and illegal poaching influenced the population size, distribution ranges and population genetic structure of several deer species during the last few centuries (Balakrishnan et al., 2003; Kumar et al., 2016; Gupta et al., 2018). As the Indian subcontinent sustains a diverse ecosystem that supported high faunal richness and diversity, but very less is known about the species diversification, evolutionary history, and how species were impacted by environmental changes during the Late Pleistocene. (Mani, 1974; Patrick et al., 2014;). The wide distribution of the northern red muntjac makes this species an excellent model to study the biogeography of Southeast Asia.

The northern red muntjac is currently listed under the “Least Concern” category in the IUCN Red List and is protected under Schedule III of the Indian Wild Life (Protection) Act, 1972. The extensive sampling of northern red muntjac from India can provide a clear depiction of population boundaries and the genetic structuring for its classification and implementing the management plan. Recently, Martin et al. (2017) investigated the geographic distribution of mtDNA lineages among red muntjac populations using museums, zoos, and opportunistically collected samples from Vietnam, Laos, and Peninsular Malaysia. As these samples belong to museums, therefore the complete mitogenome generated from this study contains lots of ambiguous nucleotides in most of the samples. Further, Martin et al. (2017) suggested extensive sampling to unveil taxonomic uncertainties of red muntjac with an analysis of nuclear data to examine barriers to gene flow. Groves et al. (2011) have suspected the presence of a Northwest lineage from Northwestern, Central India, and Myanmar. To test this hypothesis, we aimed to investigate the phylogeography and molecular ecology of red muntjac from India. We used complete mitogenome sequences data to gain insights into the evolutionary history and phylogeographic pattern; and microsatellite loci to address the population genetic structure, genetic variation, and differentiation. Based on these estimates, we can understand the speciation events of mammals in Southeast Asia impacted by biogeographical changes during the Late Pleistocene.

## MATERIALS AND METHODS

### Sample collection and DNA extraction

We used 42 archival samples of northern red muntjac from a different region of India including northwest region (NWI=18), mainland (localities of north and central) (MLI=14), northeastern (NEI=3), and southern India (SI=5) and Andaman & Nicobar Islands (AI=2) (Table 1 and Fig. 1). Genomic DNA (gDNA) was extracted from tissue and hair samples using modified DNeasy Blood & Tissue kit (Qiagen, Hilden, Germany) protocol, whereas, for antlers and bone samples, we followed Gu-HCl based silica binding method (Gupta et al., 2013). These samples were collected from dead animals from the known localities by the local Forest Department of India and sent to Wildlife Forensic and Conservation Genetic Cell, Wildlife Institute of India, Dehradun. Since the samples were collected from the dead animals, Animal Ethics Committee approval was not required for this study. The authors confirmed that all experiments were performed in accordance with relevant guidelines and regulations.

**Table 1.** Samples details used for genetic analysis of red muntjac from India. N represent the sample size.

| Origin | Status | mtDNA |  | Microsatellite |  |
| --- | --- | --- | --- | --- | --- |
|  |  | n | Types | n | Types |
| Northwest India | wild | 5 | Tissue | 18 | Tissue (15), Hairs (3) |
| Central India | wild | 3 | Tissue | 14 | Tissue (7), Antler (3), Hairs (4) |
| Northeastern India | wild | 2 | Tissue | 3 | Tissue (2), Bones (1) |
| Southern India | wild | 4 | Tissue | 5 | Tissue |
| Andaman & Nicobar Islands | wild | 2 | Tissue | 2 | Tissue |

**Figure 1.**
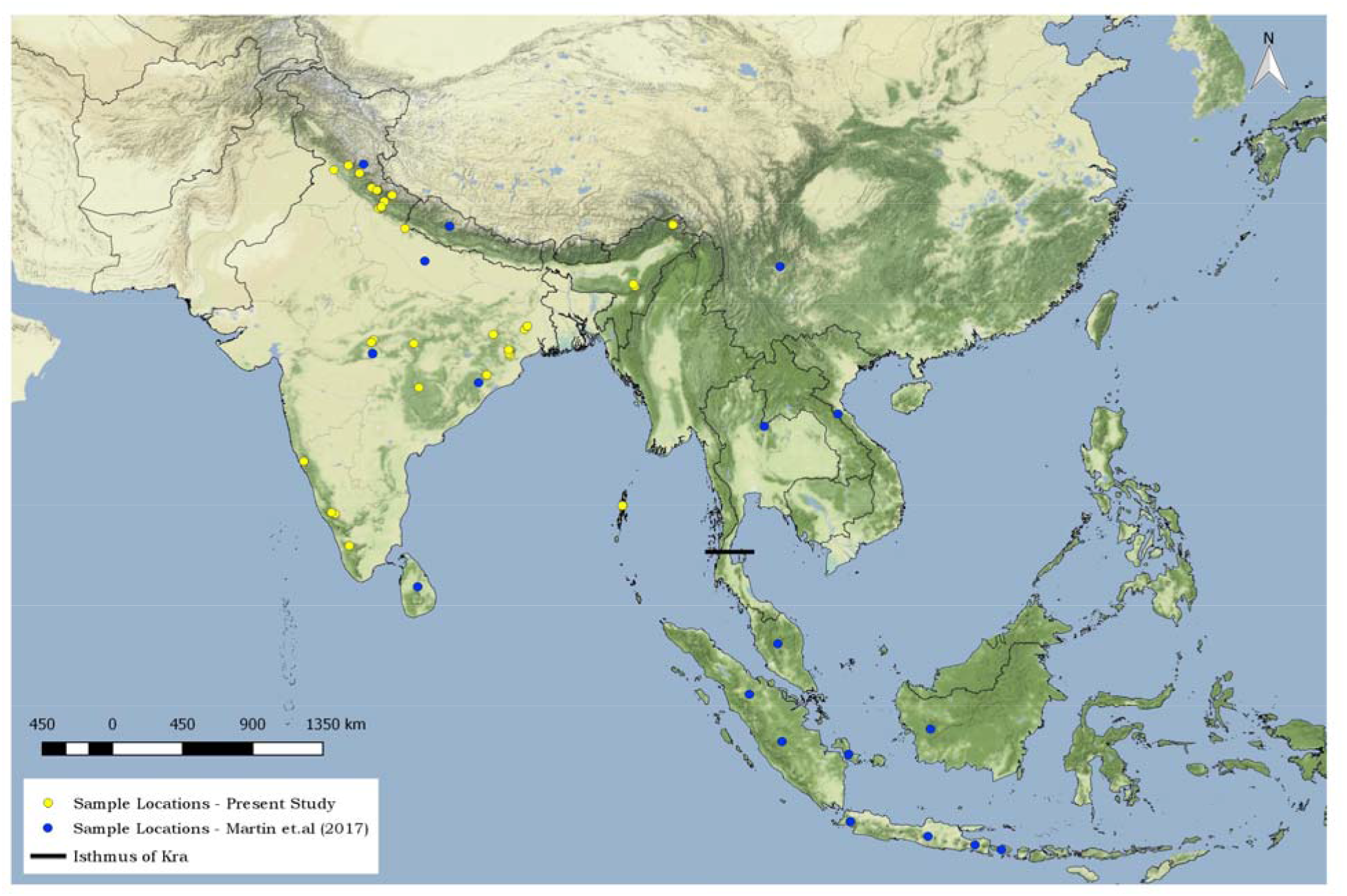
Map depicting the sampling sites of red muntjacs. Yellow dots represent samples collected for the present study; blue dots indicate the sampling location used in Martins et al., 2017. The backline indicates the position of the Isthmus of Kra.

### PCR amplification of complete mitogenome and sequencing

Polymerase chain reactions (PCRs) amplifications were carried out in 20μl volumes containing 10–20 ng of extracted genomic DNA containing, 1× PCR buffer (Applied Biosystem), 2.5mM MgCl_2_, 0.2 mM of each dNTP, 5 pmol of each primer, and 5 units of Taq DNA polymerase (Thermo Scientific). We performed PCR amplification using 23 overlapping fragments of complete mtDNA (Hassain et al. 2009). Besides, we included the fragments of complete Cytochrome *b* (Gupta et al., 2014) and Cytochrome C oxidase-*I* gene (Kumar et al., 2017) to increase the overlapping. PCR cycles for DNA amplification were 95°C for 5 min; followed by 35 cycles of 95°C for 40 sec. (denaturation), 54-56°C (annealing) for 40 sec, 72°C for 50 sec. (extension) and a final extension of 72°C for 15 min. The efficiency and reliability of PCR reactions were monitored by using control reactions. The PCR products were electrophoresed on 2% agarose gel and visualized under UV light in the presence of ethidium bromide dye. The amplified PCR products were treated with Exonuclease-I and Shrimp alkaine phosphatase (USB, Cleveland, OH) for 15 min. each at 37°C and 80°C, respectively, to remove any residual primer. The purified Amplicons were then sequenced bidirectionally using BigDye Terminator 3.1 on an automated Genetic Analyzer 3500XL (Applied Biosystems, Carlsbad, CA, USA).

### Microsatellite loci amplification and genotyping

Nine cross-species microsatellite loci: INRA001 (Vaiman et al. 1992), Ca18 (Gaur et al. 2003), BM6506 (Bishop et al. 1994), RT1, RT27, NVHRT48 (Poetsch et al. 2001), CelJP27 (Marshall et al. 1998), and T156, T193 (Jones et al. 2002) were selected for population genetic analysis of the red muntjac. Three sets of the multiplex set were carefully assembled based on molecular size and labeled fluorescent dyes of loci. Multiplex amplification was carried out in 10 μl reaction volumes containing 5 μl of QIAGEN Multiplex PCR Buffer Mix (QIAGEN Inc.), 0.2 μM labeled forward primer (Applied Biosystems), 0.2 μM unlabeled reverse primer, and 20–100 ng of the template DNA. PCR cycles for loci amplification were 95°C for 15 min; followed by 35 cycles of 95°C for 45 sec. (denaturation), 55°C (annealing) for 1 min, 72°C for 1 min. (extension) and a final extension of 60 °C for 30 min. Alleles were resolved in an ABI 3500XL Genetic Analyzer (Applied Biosystems) using the LIZ 500 Size Standard (Applied Biosystems) and analyzed using GeneMaker ver 2.7.4 (Hulce et al., 2011).

### Data Analysis

A total of 16 complete mitogenome sequences of red muntjac from five different localities of India were generated (Table 1). Sequences obtained from forward and reverse direction were edited and assembled using SEQUENCHER®version 4.9 (Gene Codes Corporation, Ann Arbor, MI, USA). The annotation of complete mitogenome was done using Mitos WebServer (Bernt et al., 2013) and MitoFish (Iwasaki et al., 2013). Careful manual annotation was also conducted by sequence alignment with their related homologs sequences or species for ensuring the gene boundaries. Additionally, 36 sequences that consisted of northern (n=17); southern (n=17) and Sri Lankan (n=2) muntjac were included from GenBank to cover wide distribution ranges for better insight (Fig. 1). The dataset of 52 sequences of red muntjac were aligned using the CLUSTAL X 1.8 multiple alignment program (Thompson et al., 1997) and alignments were checked by visual inspection. DnaSP v 5 (Librado and Rozas, 2009) was used to estimate the haplotype (h) and nucleotide (π) diversity. A phylogenetic analysis was conducted in BEAST ver 1.7 (Drummond et al., 2012). For a better insight into the phylogenetic relationships, *Bos javanicus* (JN632606) was used as an out-group. The resulting phylogenetic trees were visualized in FigTree v1.4.0 (http://tree.bio.ed.ac.uk/software/figtree/). The spatial distribution of haplotypes was visualized by a median-joining network, which was created using the PopART software (Leigh and Bryant, 2015). The genetic distance between lineages was calculated based on Tamura-3 parameter with a discrete Gamma distribution (TN92+G) with the lowest BIC score value implemented in MEGA X (Kumar et al., 2017).

### Estimating divergence dating

To estimate divergence times of red muntjac clades, we inferred genealogies using a relaxed lognormal clock model in BEAST ver 1.7 (Drummond et al., 2012). We performed the dating estimates using fifteen sequences downloaded from NCBI, including the species of Cervidae and Muntiacini. i.e., the Chital (*A. axis*, JN632599), Swamp deer (*R. duvaucelii*, NC020743), Red deer (*C. elaphus*, AB245427), European Roe deer (*C. capreolus*, KJ681491), Fallow deer (*D. dama*, JN632629), Water deer (*H. inermis*, NC011821), Mule deer (*O. hemionus*, JN632670), Hog deer (*A. porcinus*, MH443786), Formosan sambar (*R.u.swinhoei*, DQ989636), Tufted deer (*E. cephalophus*, DQ873526.), Chinese muntjac (*M. reevesi*, NC_008491), Giant muntjac (*M. vuquangensis*, FJ705435), Putao muntjac (*M. puhoatensis*, MF737190) and Black muntjac (*M. crinifrons*, NC_004577) and the sequence of one species of the family Bovidae, i.e., the Banteng (*B. javanicus*, FJ997262). The TMRCA of bovids and cervids was set at a calibration point to 16.6 ± 2 million years ago (Mya) with a normal prior distribution (Martin et al.,2017, Bibi et al.,2013). We used a Yule-type speciation model and the HKY + I + G substitution rate model for tree reconstruction. We conducted two independent analyses, using MCMC lengths of 10 million generations, logging every 1000 generation. All the runs were evaluated in Tracer v. 1.6. The final phylogenetic tree was visualized in FigTree v.1.4.2 (http://tree.bio.ed.ac.uk/software/figtree/).

### Microsatellite analysis

A total of nine polymorphic loci were used to analyze the 42 samples of red muntjac for population genetic studies. The CERVUS ver 3.0.6 program (Kalinowski et al., 2007) was used to estimate the polymorphic information content (PIC), the number of alleles per locus, the observed (H_o_) and, expected (H_E_) heterozygosity. The allelic richness (Ar) and mean inbreeding coefficient (F_*IS*_) (Weir et al., 1984) was estimated using FSTAT ver 2.9.3 (Goudet et al., 1995). All the loci were checked for under HWE in GenAlEx v6.5 (Peakall and Smouse, 2012). Factor correlation analysis (FCA) was performed using the GENETIX 4.05 software package (Belkhir et al., 2004). CONVERT 1.31 (Glaubitz et al., 2007) was used to convert the required input file format.

To determine the genetic structure of population, the DAPC method was implemented using ADEGENET package in R (Jombart et al.,2005). DAPC is a multivariate and model-free approach that maximizes the genetic differentiation between groups with unknown prior clusters, thus improving the discrimination of populations. This analysis does not require a population to be in HWE. The pairwise *F*_ST_ values (gene flow) among the populations were calculated using GenAlEx v6.5 (Peakall and Smouse, 2012).

### Spatial genetic analysis

The correlation between the pairwise genetic and geographic distances was performed to detect the pattern of isolation by distance between the disjointed areas, according to Mantel’s test implemented in Alleles in Space 1.0 (Miller et al., 2005).

## RESULTS

### Phylogeographical analyses

The generated 16 mitogenomes sequences of red muntjac were deposited in GenBank. To depict the phylogeographical relationships, we included four sequences of *M. vaginalis* from Singh et al., (2018) and 32 sequences of northern, southern, and Srilankan muntjac from Martin et al. (2017). All complete mitogenomes sequences represented individual haplotypes, indicating maternal unrelatedness of all red muntjac samples.

The Bayesian consensus tree showed that all the sequences of red muntjac were clustered into four major clades (Figure 2). Clade-*I* consists of the sequences of northern red muntjac lineages (*M. vaginalis*), which comprises individuals ranging from north to central India, eastern to north-east India, Nepal, Southeast Asia, and Andaman & Nicobar Islands. Clade-*II* comprises individuals from the Northwestern part of India (i.e., Uttarakhand, Punjab, and Himanchal Pradesh). Clade-*III* comprises the sequences originated from Sunda (*M. muntjak*), mainly from Sumatra, Malay Peninsula, Lombok, Borneo, Java, and Bali’s Islands. Clade-IV consists of individuals from Western Ghats of Southern Indian and Sri Lankan red muntjac (*M. malabaricus*). The median-joining (MJ) network of all recognized sequences from India, South East Asia Sundaland, and Sri Lanka strongly supported phylogenetic results. It indicated the existence of four geospatial population structuring from their distribution ranges (Fig. 3). In India, we found three evident clades: 1) comprised of Northwest India, 2) mainland populations, and 3) the Western Ghats. Interestingly, the phylogenetic and MJ analysis indicated the existence of two sub-groups in the Sunda of Southern red muntjac. It is also noteworthy that the mainland red muntjacs from India were exhibited different genetic signature and showed structuring with respect to other mainland population that existed in Southeast Asia and Andaman & Nicobar Islands. The nucleotide diversity among red muntjac mitogenomes was estimated according to their clustering pattern. The nucleotide diversity among northern lineages was π =0.0044 (s.d.=0.0003), north-west was π =0.0016 (s.d.=0.0002), Western Ghats + Sri Lankan lineage was π =0.005 (s.d.=0.0008) and Sundaland was π =0.0109 (s.d.=0.0006). The overall nucleotide diversity among red muntjac was π =0.0203 (s.d.=0.001).

**Figure 2.**
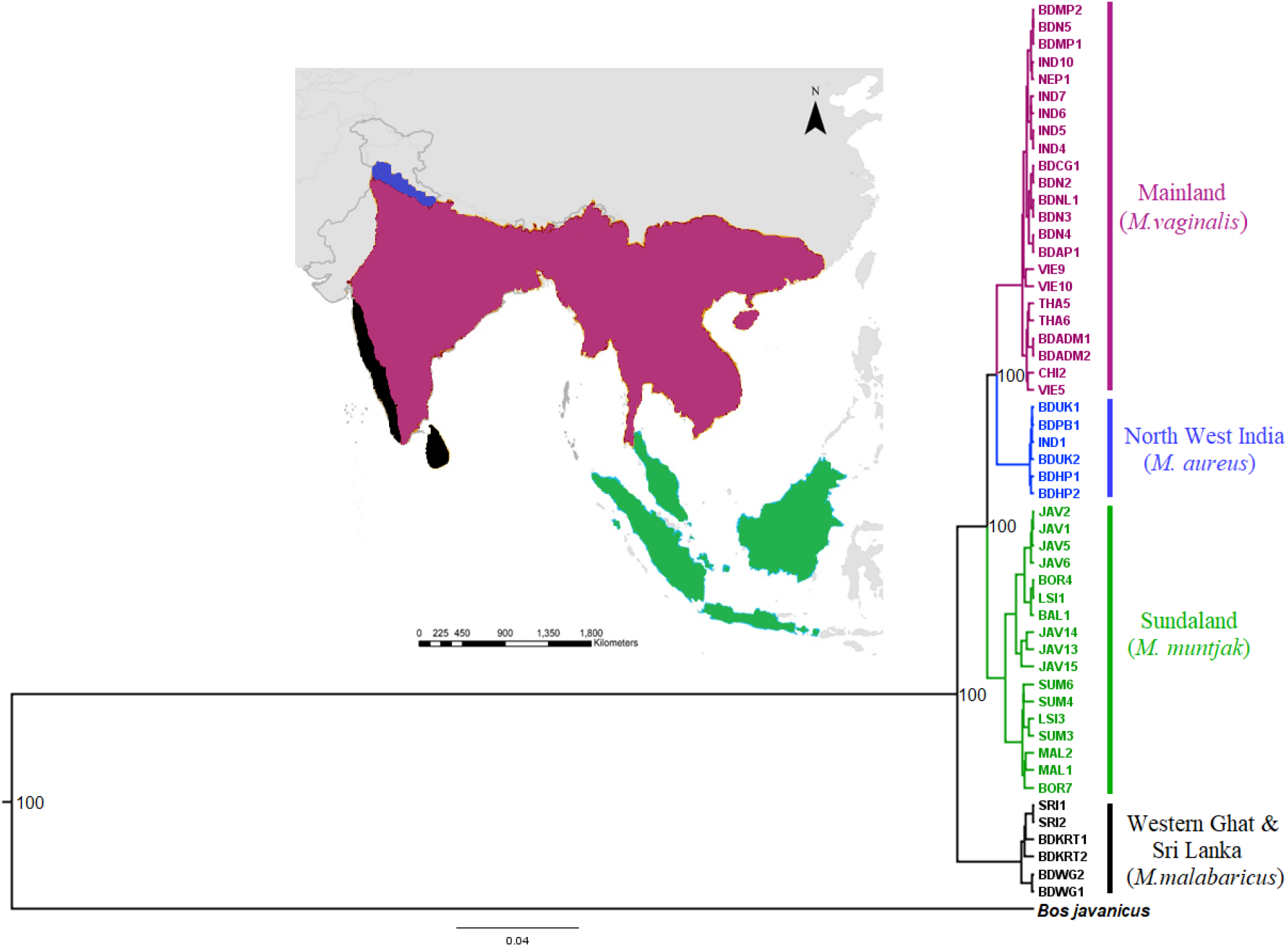
Bayesian inference (BI) phylogenetic tree for red muntjac based on complete mitochondrial DNA. Numbers on clades indicate posterior probability for the node. The distribution ranges of different lineages are represented by specific colors corresponding to colored clades in a tree topology. The *Bos javanicus* (FJ997262) was used as an outgroup.

**Figure 3.**
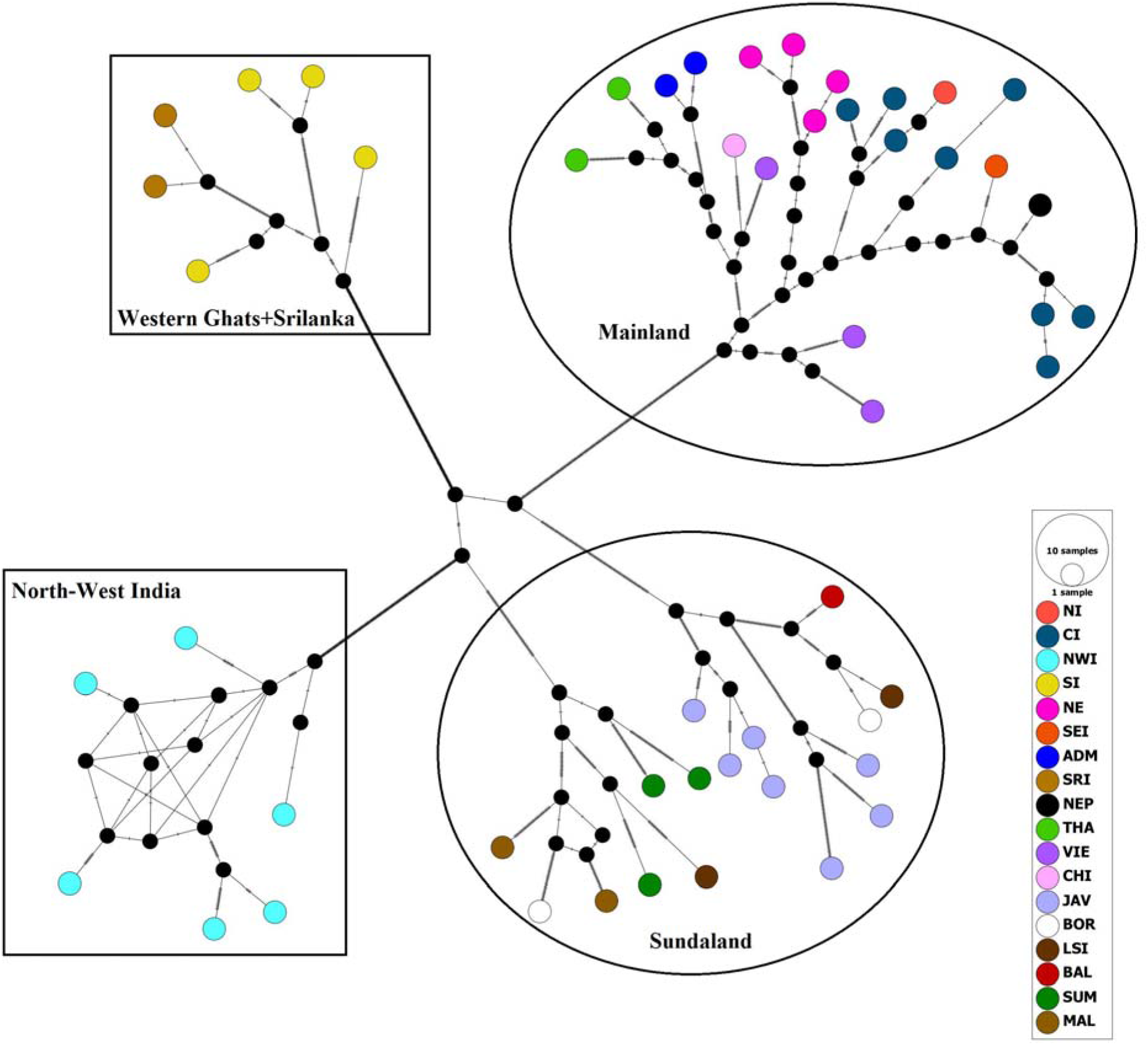
A median-joining (MJ) network of full mitogenome of red muntjac lineages. The size of each circle indicates the relative frequency of the corresponding haplotype in the whole dataset.

### Estimating genetic divergence

The complete mitogenome dataset was used to estimate the divergence time to a most recent common ancestor (TMRCA) of the Cervids and Bovids at 16.6 ± 2 million years (Myr); CI_95%_: 11.7–19.3 Myr. Our divergence results suggested that the splits between the red muntjac and black muntjac (*M. criniforns*) occurred in the Late Pliocene, around 2.68 Myr (CI_95%_: 1.77–3.81). Within red muntjac, group diversification started during Pleistocene. The *M. malabaricus* split earlier approximately 1.7 Myr (CI_95%_:1.23–2.67); after that, the clade of Southern red muntjac of Sunda split around 1.16 Myr (CI_95%_:0.75–1.68). Within northern lineages, the split between the north and northwest lineage was estimated to occur at about 0.83 Myr (CI_95%_:0.53–1.26) (Fig. 4). According to our estimates, the northwest red muntjac was a younger group among red muntjac.

**Figure 4.**
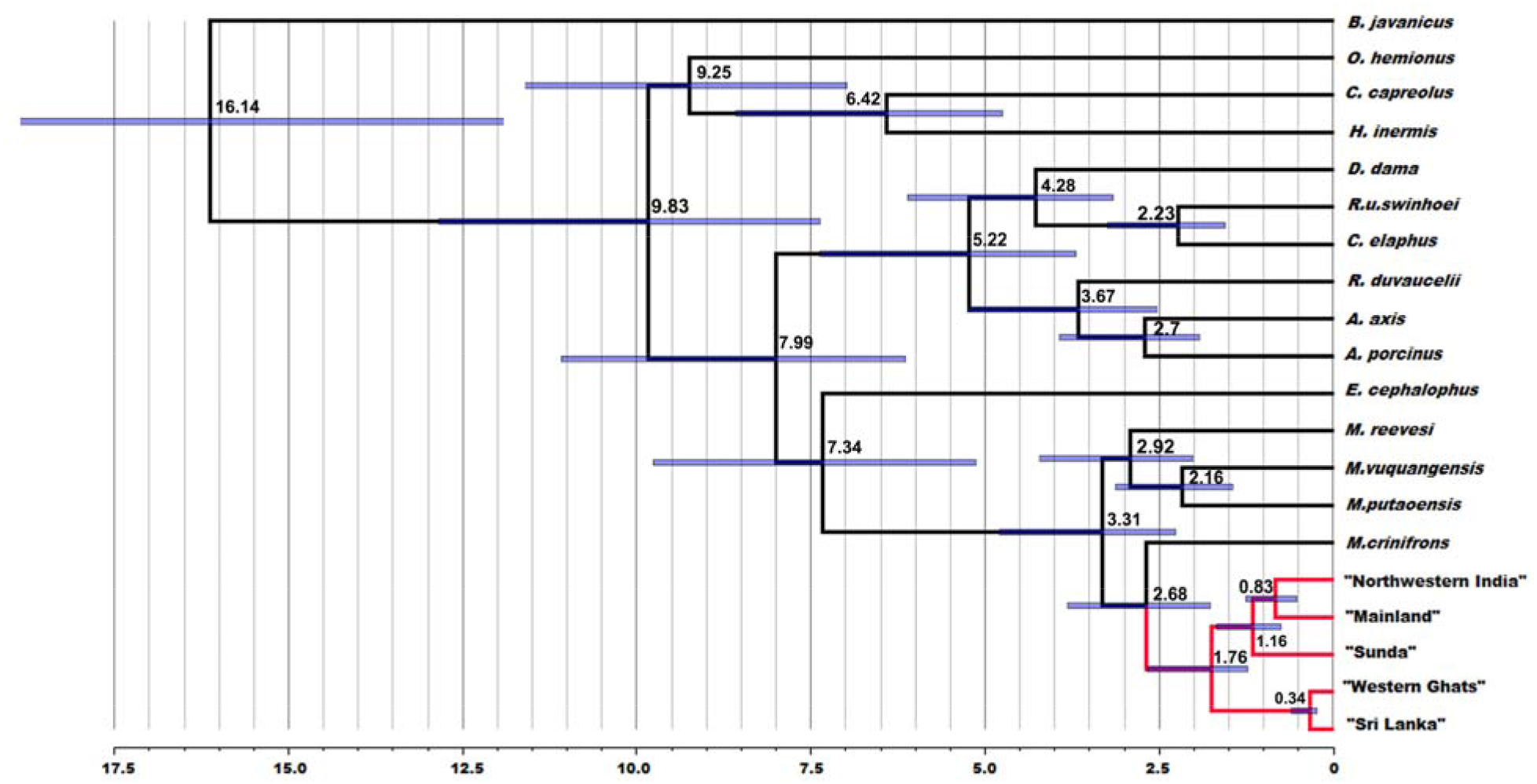
Complete mitogenome based maximum credibility tree was generated using BEAST. Molecular clocks were computed after calibrating the root age between bovids and cervids at 16.6 ± 2 Mya (CI_95%_: 11.7–19.3). The red clade shows that the split between the four lineages of the red muntjac occurred at around 2.68 Myr (CI_95%_: 1.77–3.81).

### Genetic diversity

The genetic diversity of Indian red muntjacs was calculated using nine microsatellite markers (Table 2). The selected microsatellite markers showed high polymorphic information content (PIC>0.5) with a mean value of 0.831; therefore, all used loci were found to be informative. All loci significantly deviated from Hardy-Weinberg equilibrium, and P-value was 0. No linkage disequilibrium was detected (P *>* 0.05). In the overall population, the observed allele per locus was ranging from 4 to 9, with a mean 7.00. The observed (*Ho*) and expected heterozygosity (*H*_*E*_) of northwest population *Ho*: 0.666; *H*_*E*_: 0.730, mainland population *Ho*: 0.619; *H*_*E*_: 0.823, whereas in the Western Ghats, it was *Ho*: 0.756; *H*_*E*_: 0.760 with mean 0.680 and 0.771, respectively.

**Table 2.** Summary of genetic diversity in hog deer populations of India. Key: Na = number of alleles; Ne = number of effective alleles; Ho = observed heterozygosity; HE = expected heterozygosity; PIC = Polymorphic information content. HWE ***P < 0.001; Indicative adjusted nominal level (5%) = 0.001.

| Loci | All population (n=42) |  |  |  |  |  | Lineage 1<br>North western India<br>(n=18) |  |  | Lineage 2<br>Mainland India (n=19) |  |  | Lineage 3<br>Southern India (n=5) |  |  |
| --- | --- | --- | --- | --- | --- | --- | --- | --- | --- | --- | --- | --- | --- | --- | --- |
|  | Size Range | Na | Ne | Ho | HE | PIC | Na | Ho | HE | Na | Ho | HE | Na | Ho | HE |
| BM6506 | 184-200 | 8 | 5.522 | 0.759 | 0.811 | 0.842 | 8 | 0.833 | 0.752 | 11 | 0.444 | 0.841 | 7 | 1.000 | 0.840 |
| INRA001 | 193-225 | 7 | 4.280 | 0.455 | 0.730 | 0.817 | 5 | 0.438 | 0.592 | 10 | 0.526 | 0.839 | 6 | 0.400 | 0.760 |
| RT1 | 205-231 | 6 | 5.226 | 0.778 | 0.794 | 0.811 | 5 | 0.778 | 0.708 | 7 | 0.556 | 0.833 | 8 | 1.000 | 0.840 |
| T193 | 165-189 | 6 | 4.584 | 0.789 | 0.780 | 0.841 | 8 | 0.778 | 0.778 | 6 | 0.789 | 0.803 | 6 | 0.800 | 0.760 |
| RT27 | 135-155 | 9 | 5.611 | 0.772 | 0.755 | 0.899 | 10 | 0.778 | 0.787 | 14 | 0.737 | 0.898 | 4 | 0.800 | 0.580 |
| AY302223 | 195-223 | 7 | 6.078 | 0.671 | 0.831 | 0.869 | 6 | 0.833 | 0.790 | 10 | 0.579 | 0.861 | 7 | 0.600 | 0.840 |
| NVHRT48 | 105-115 | 9 | 5.823 | 0.742 | 0.820 | 0.882 | 10 | 0.778 | 0.773 | 11 | 0.647 | 0.867 | 6 | 0.800 | 0.820 |
| CelJP27 | 176-196 | 4 | 2.828 | 0.529 | 0.620 | 0.701 | 4 | 0.222 | 0.569 | 7 | 0.765 | 0.749 | 3 | 0.600 | 0.540 |
| T156 | 131-227 | 9 | 5.438 | 0.627 | 0.799 | 0.825 | 8 | 0.556 | 0.824 | 11 | 0.526 | 0.713 | 8 | 0.800 | 0.860 |
| Mean |  | 7 | 5.043 | 0.680 | 0.771 | 0.831 | 7.111 | 0.666 | 0.730 | 9.667 | 0.619 | 0.823 | 6.111 | 0.756 | 0.760 |
| S.E. |  | 0.49 | 0.349 | 0.035 | 0.019 |  | 0.735 | 0.071 | 0.030 | 0.850 | 0.041 | 0.020 | 0.564 | 0.065 | 0.040 |

### Population genetic structure and genetic differentiation

We used DAPC to analyze the genetic structure of the red muntjac population from India. We found that there were three genetic clusters: Cluster-I northwest India, Cluster-II mainland, and Cluster-III Western Ghats of Southern India (Fig. 5 and Fig. 6). The factor correlation analysis (FCA) differentiated the population of northwest India, mainland, and Western Ghats (Fig. 7). These clusters were highly supported by mitogenome data.

**Figure 5.**
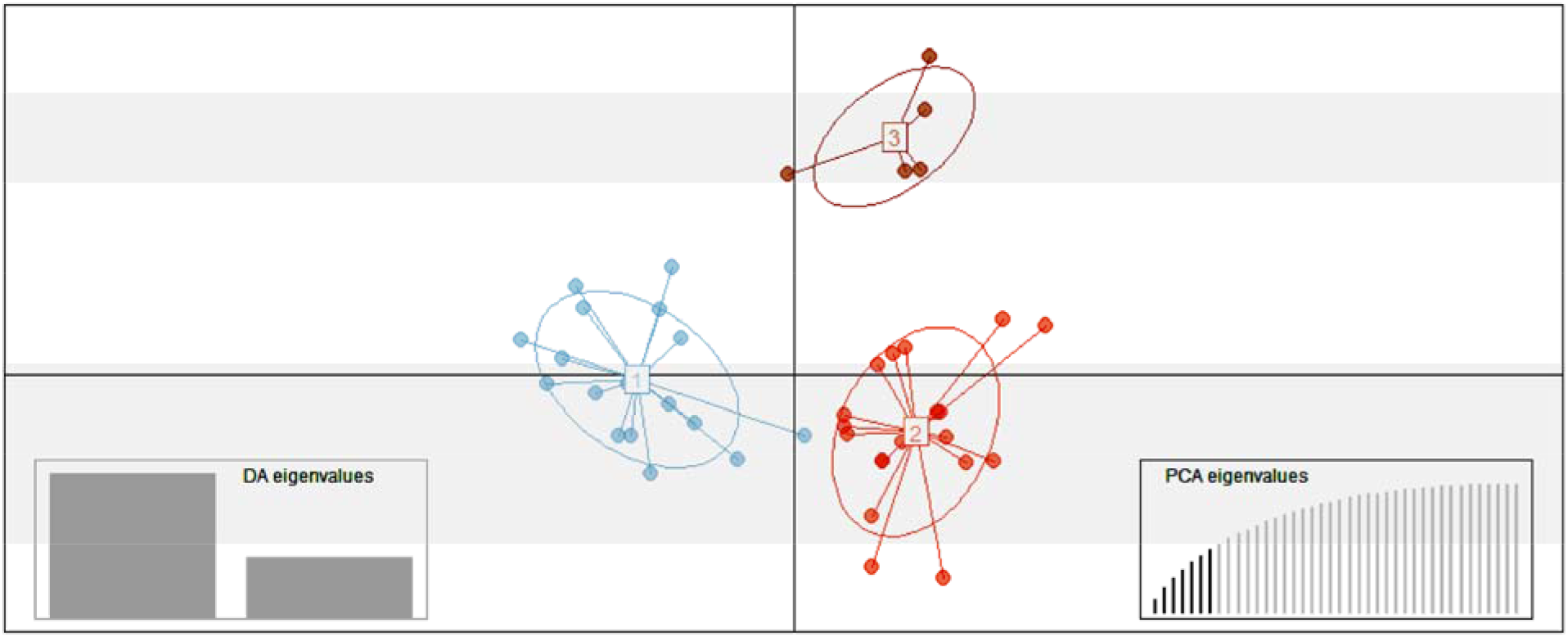
Scatterplots of DAPC based on microsatellite loci for three red muntjac populations. Dots represent individuals. DAPC was carried out using a hierarchical islands model and shown by different colors and inertia ellipses. The DA and PCA eigenvalues of the analysis are displayed in the inset.

**Figure 6.**
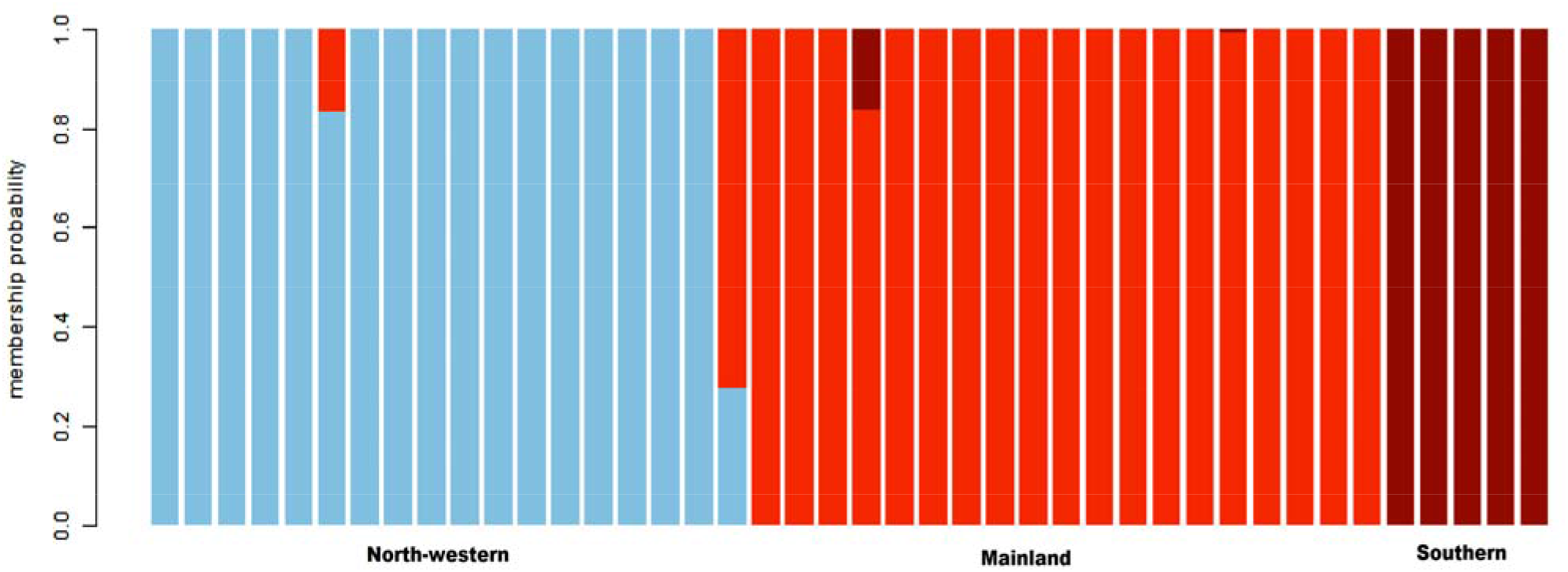
Results of bar plot showing genetic clustering implemented in Discriminant Analysis of Principal Components (DAPC) based on microsatellite data. Each column along the X-axis represents one red muntjac individual. The Y-axis represents the assignment of the membership probability of each individual.

**Figure 7:**
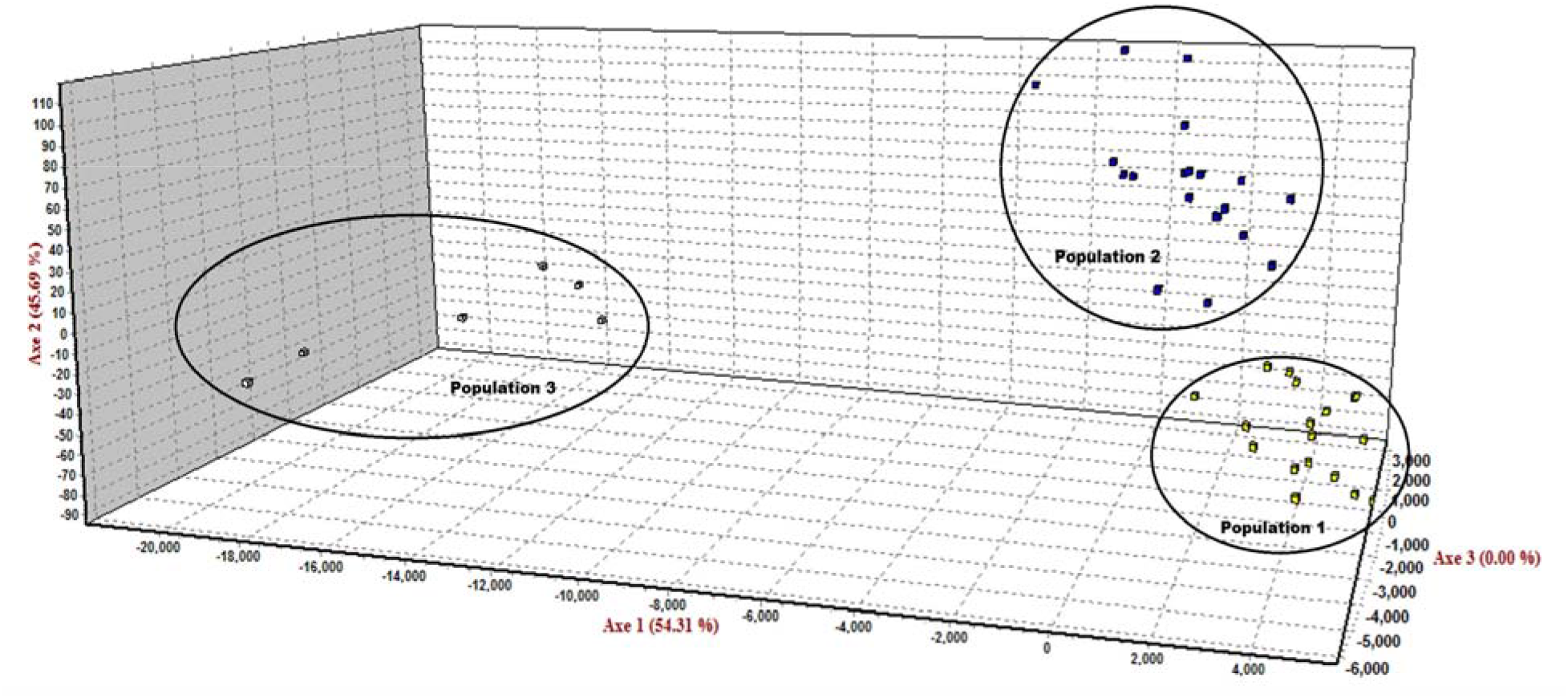
Results of factor correlation analysis (FCA) using microsatellite markers, indicating three major clusters in the Red muntjac population in India.

The analysis based on pairwise *F*_*ST*_ for red muntjac based on complete mitogenome demonstrated significant genetic differentiation from the mainland to the northwest population (0.0208), and Western Ghat+Srilanka (0.0396). The genetic differentiation from North-west and Mainland to the Sunda population is almost similar (0.026), whereas a high level of differentiation was observed between Sunda and Western Ghat+Srilanka (0.039). A similar pattern was observed with microsatellite analysis where the observed genetic differentiation between northwest and the mainland was 0.061, and Western Ghat is 0.105, whereas mainland to Western Ghat was 0.080 (Table 3). We also calculated genetic differentiation with other Muntiacus species where we found a comparatively low level of genetic differentiation between red muntjac lineages with *M. criniforns* (0.056 to 0.057) than the other species such as *M. reevesi* (0.065 to 0.068)*, M. vuquangensis* (0.060 to 0.068) and *M. putaoensis* (0.067 to 0.069). The spatial genetic analysis revealed a significant correlation between the pairwise genetic distances among geographical subsets and geographical distances (Mantel test, *rM* = 0.513; *P* = 0.0009, Fig.8). This pattern of isolation by distance (IBD) was strongly influenced by the genetic differentiation and the major geographical distance between the red muntjac lineages.

**Table 3.** Genetic differentiation among red muntjac and other Muntiacus species. The pairwise *FST* values based on the complete mitogenome and microsatellite loci (in bold) are shown above and above the diagonal, respectively.

| Species | 1 | 2 | 3 | 4 | 5 | 6 | 7 | 8 |
| --- | --- | --- | --- | --- | --- | --- | --- | --- |
| *M.vaginalis* | - | **0.061** | **0.080** |  |  |  |  |  |
| *M.aureus* | 0.0208 | - | **0.105** |  |  |  |  |  |
| *M. malabaricus* | 0.0396 | 0.0401 | - |  |  |  |  |  |
| *M. muntjak* | 0.0269 | 0.0264 | 0.0399 | - |  |  |  |  |
| *M. crinifrons* | 0.0557 | 0.0572 | 0.0562 | 0.0571 | - |  |  |  |
| *M. reevesi* | 0.0668 | 0.0657 | 0.0682 | 0.0685 | 0.0651 | - |  |  |
| *M. vuquangensis* | 0.0679 | 0.0683 | 0.0672 | 0.0681 | 0.0665 | 0.0606 | - |  |
| *M. putaoensis* | 0.0693 | 0.0690 | 0.0676 | 0.0695 | 0.0692 | 0.0624 | 0.0486 | - |

**Figure 8:**
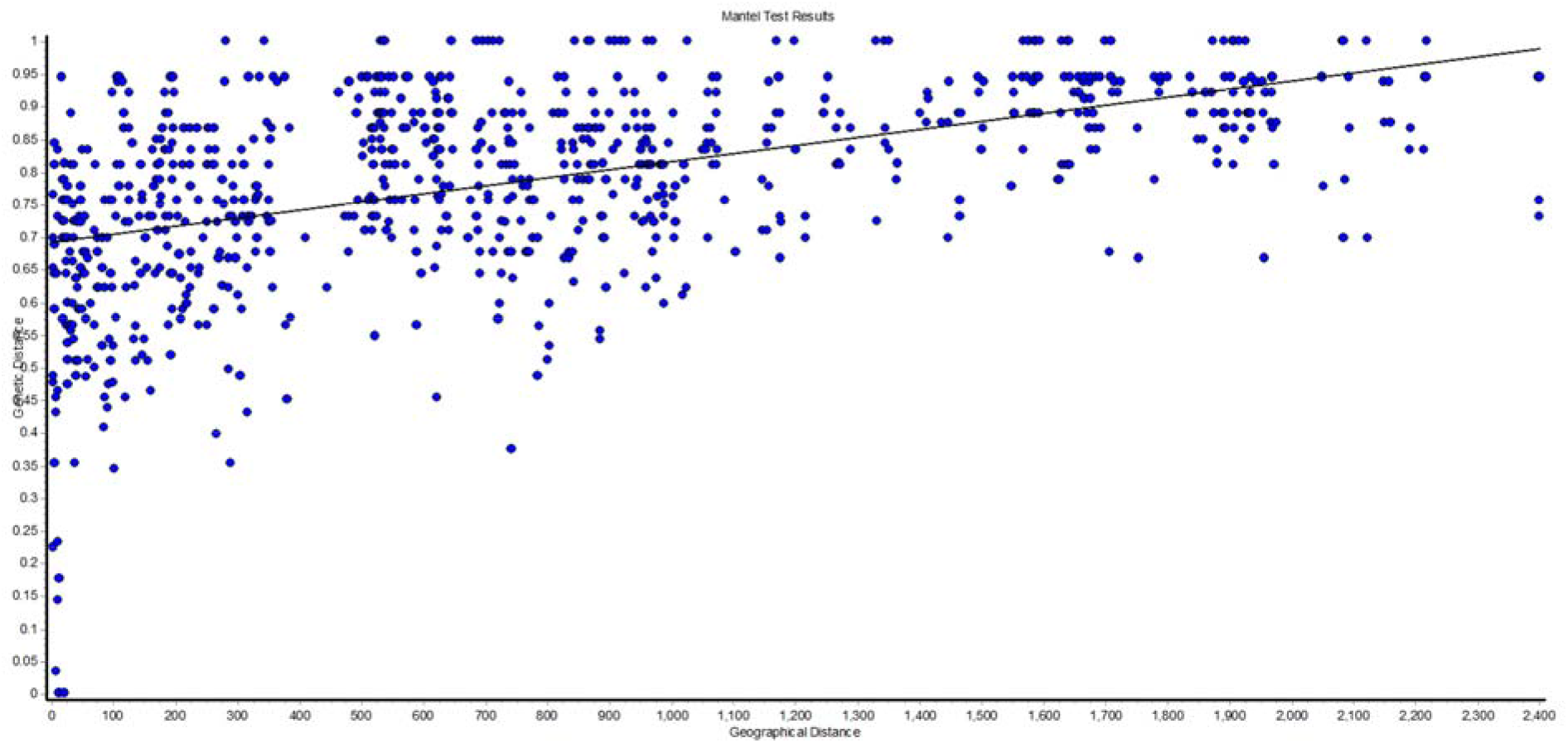
Correlation of genetic and geographical distance in kilometer between red muntjac population using microsatellite data (Mantel test, *rM* = 0.513; *P* = 0.0009).

## DISCUSSION

This study is a pioneer work that brings together the extensive analyses of complete mitochondrial and microsatellite loci variation to understand the Quaternary climatic fluctuations and geological events on the probable influence on the demographic pattern and population genetic structure among the red muntjac groups. Phylogeographic studies of red muntjac groups showed a clear division between northern and southern lineages, indicating the physical barriers to gene flow that have existed as a result of extreme dry climatic conditions caused by global ice advances (Martin et al., 2017). As the taxonomy of muntjacs are considerably fragile and has been in the debate (Groves et al., 2011), and research is still ongoing to resolve the phylogenetic complexities to answer the existence of the exact number of the lineages. The population genetic studies on red muntjacs will act as dominant tools for understanding the lineages diversification, genetic structuring, and diversity, which has resulted in the development of appropriate conservation and management strategies for this enigmatic species.

### Phylogeographic analysis and population structure

Present studies of genetic structure and differentiation among red muntjac populations exhibit the existence of four genetically distinct lineages from its geographical distribution range in South and Southeast Asia. Previously, Martin et al, (2017) reported three mitochondrial lineages: The Sri-Lankan red muntjac (*M. malabaricus*), Northern red muntjac (*M. vaginalis*) and Southern red muntjac (*M. muntjak*). Based on our mitogenome and microsatellite data, we identified the presence of a new lineage named Himalayan red muntjac (*M. (m.) aureus*) of red muntjac from Northwest part of India. Our mitogenome data estimated that the Northwest lineages and mainland lineage split during the late Pleistocene approximately 0.83 Myr (CI_95%_:0.53–1.26), which is a younger lineage among red muntjac. Himalayan red muntjac is inhabited in foothills of Himalayan high elevated Northwestern part of Indian states of Uttarakhand, Himachal Pradesh, and Punjab. The distribution range of Northwest lineage was also suspected to occur from Northwestern as well as Central India and Myanmar (Groves et al. 2011). The presence of distinct genetic lineages from the Northwest Indian region was also speculated by Martin et al. (2017) based on one sequence from Himachal Pradesh Province that formed a distinct placement. However, due to the unavailability of the appropriate sample size, the explicit depiction of this lineage was lacking. Hence, this study proven the previous concept/hypothesis and provided molecular evidence for the confirmation of Northwest lineage from India. The rise of anthropogenic activities in Late Quaternary become a key factor that changed the pattern of global biodiversity (Turvey et al., 2016). The upliftment of Qinghai-Tibetan Plateau (QTP) plays a significant role in faunal and floral diversification and evolution in Himalayan ranges (Favre et al., 2015). The upliftment of the Himalayas followed by the Plio-Pleistocene glaciations has led to the evolution of high altitudinal elements shaping biogeographic evolution in the Indian Himalayan region (Mani et al., 1974). The Himalayan region enabled allopatric speciation through geographic isolation as well as adaptive diversification across altitudinal gradients (Doebeli & Dieckmann, 2003; Korner, 2007). Such diversification was also driven by pre-adapted lineages immigrating and undergoing subsequent speciation (Johansson et al., 2007; Price et al., 2014). After that, rapid civilization in the Indo-Gangetic plain of North India caused extensive destruction of natural habitat, which altered the ecology, distributional pattern, and range of plants and animals (Mani, 1974).

Based on our phylogenetic study, we could not find any genetic signature of the Northwestern lineage of red muntjac in the central part of India. In congruent to their distribution pattern, all red muntjac population sustains a high mtDNA and microsatellite diversity with large divergence among them. Our result indicated that Western Ghats lineage is genetically more diverse than the mainland, northwest, and Sundaland populations. The phylogenetic result showed that the Srilankan red muntjac (*M. malabaricus*) is the most primitive lineage of red muntjac that is also distributed until the Western Ghats in India. The genetic divergence result suggested that Srilankan red muntjac split from the mainland lineage during Pleistocene around 1.7 Mya, when the climatic condition was quite complex in India (Martin et al., 2017). Despite the present biogeographic separation between Southern India and Sri Lanka, both populations are genetically similar at the mitogenomic level. A similar genetic relationship was also observed in *Paradoxurus* (palm civets), where Southern India and Sri Lanka clustered with each other (Patou et al., 2010). The common origin of Western Ghat, India and Srilankan lineages might be the results of historical connectivity between these two landscapes, and after that, the changes in sea levels probably led to the isolation that might have resulted in the existence of local endemism (Patou et al., 2010; Bossuyt et al., 2004; Manamendra-Arachchi et al., 2005). In India, the discontinuity of Western Ghat to mainland population resulted due to unfavorable habitat conditions that culminated in isolated patches forming the refugia population inhabited in Western Ghats biodiversity hotspot (Karanth et al. 2003). The barrier formed by the central dry zone of India restricted the geneflow between the Western Ghats and the rest of the Indian population of red muntjac. The restriction in species distribution is also probably due to pronounced climate fluctuation in last glacial maxima that caused contraction of rain forest; therefore, forest-dwelling species have to restrict themselves to the only available habitat of high elevation areas (Meijaard & Groves, 2006, Patou et al.,2010).

The mainland red muntjac (*M. vagianalis*) distributed in a larger landscape of India (i.e, Chhattisgarh, Odisha, West Bengal, North East India, and Andaman & Nicobar Islands) and Indo-Chinese region (Vietnam, China and Thailand). The Andaman & Nicobar Islands samples were genetically closer to the population of Indo-Chinese red muntjac. Due to unsuitable conditions in most of the areas, the mainland population limited to in a small humid forest, after the interglaciation forest prevailed into the drier areas helped former distribution to reoccupy and prolongation of ranges by red muntjacs. Southern red muntjac inhabited in Sudanic region (Java, Sumatra, Bali and Borneo) and lineage diversification was well described by Martin et al, (2017). The major transition zone between the Indochinese and Sundaic zoogeographic subregion is Isthmus of Kra located on the Malay/Thai Peninsula that might act as a possible barrier preventing gene flow between populations (Meijaard 2009; Woodruff and Turner, 2009). It is also speculated that there was no geophysical barrier in this area but suggested repeated rapid sea-level changes resulted in the isolation of species in a particular region (Woodruff and Turner, 2009). Southern red muntjac split around 1.1 Mya with the Indochinese mainland population. This divergence estimation is congruent with the lineages diversification in Amphibians (Inger and Voris 2001); Birds (Hughes et al. 2003); Mammals (Woodruff and Turner 2009); bats (Hughes et al. 2011) and Leopard cat (Patel et al.,2017) that occurred in Indochinese and Sundaic region. Faunal diversification between the Indochinese and Sundaic region might be the results of fluctuation in Indian summer monsoon (Zhisheng et al., 2011), due to rise in sea level (Miller et al., 2005; Woodruff & Turner 2009).

In conclusion, the genus *Muntiacus* is a good model for the study of the identification of criptic lineages and biogeography of Asia. This study suggested that there is a need for a taxonomic revision within red muntjac group and shown that Himalayan red muntjac (*M. (m.) aureus*) was recently separated from *M. vaginalis* and it should be managed as an evolutionary significant unit (ESUs). There is also a need to assess the conservation status under IUCN Red List category. Identifying concrete geographic limits of red muntjac can be achieved by rigorous and robust sampling in Southeast Asia. Hence, we suggest further in-depth research based on both mitochondrial and microsatellite markers to address the unclear status of red muntjacs from the Malay Peninsula. This study contributes to the understanding of the large-scale drivers of species and provides new insight into the linkage of environmental changes on the distribution and niche dynamics.

## ACKNOWLEDGEMENTS

This study was funded by the Wildlife Institute of India (WII). We acknowledge the support provided by the Director and Dean, WII. This study was supported by the infrastructure of WFCG Cell’s WII. We thank the State Forest Departments of Uttarakhand, Himachal Pradesh, Uttar Pradesh, Madhya Pradesh, Chhattisgarh, Jharkahnd, Odisha, West Bengal, Kernataka, Tamil Nadu, Nagaland, Andaman & Nicobar Islands, Assam, Arunachal Pradesh and Goa States for sending the samples.

## DATA ACCESSIBILITY

Sequence data would be available on NCBI GenBank.

## AUTHOR CONTRIBUTIONS

S.K.G. conceived the ideas; B.S. and A.K. generated the data and produced the DNA sequences; B.S., A.K. and S.K.G. analyzed the data; B.S. and A.K. led the writing; S.K.G. and V.P.U. supervised the study and writing of this paper. All authors read and approved the final version of the paper.

